# Proteoglycan Clusters as a Site of Coordinated, Multi-Dendritic Plasticity

**DOI:** 10.1101/2021.10.04.462691

**Authors:** Gabriele Chelini, Peter Durning, Sinead O’Donovan, Torsten Klengel, Luigi Balasco, Cristina Berciu, Anne Boyer-Boiteau, Yuri Bozzi, Robert McCullumsmith, Kerry J. Ressler, Sabina Berretta

**Author notes:** Correspondence to: Sabina Berretta, M.D., MRC3, Mailstop 149, 115 Mill St., Belmont MA 02478, USA.

## Abstract

Experience-dependent learning depends on synaptic plasticity. While plasticity in individual synapses has been extensively investigated, the mechanisms underlying coordinated changes across sets of synapses on multiple dendrites, likely needed to encode effective adaptations to a salient stimulus, are not well understood. The extracellular matrix is uniquely well suited to fulfill this function, as rapid glia-driven remodeling of its local composition powerfully impact synaptic plasticity. We show that extracellular matrix microenvironments, named CS6 clusters, dynamically form around several dendrites in response to sensory stimuli in coincidence to stimulus-driven synaptic plasticity. CS6 clusters, formed by glia-dependent secretion of extracellular matrix components surrounding sets of adjacent dendrites, may represent a novel structure supporting coordinated synaptic plasticity.

**One Sentence Summary:** Extracellular matrix clusters form microenvironments for coordinated multi-dendrite synaptic plasticity.

## Main Text

A critical challenge in neurobiology is to understand how neuronal activity refines local brain circuitry, resulting in experience-dependent learning. Converging evidence suggests that coordinated activity of adjacent synapses represents an efficient mechanism for acquisition and storage of novel information (*1-5*). Investigations into this mechanism, focused on dendritic spines, provide compelling evidence for local coordination of synaptic remodeling (*3, 4*). These studies show that the strength of a group of neighboring synapses on a dendrite segment increases within 90 minutes from a sensory stimulus however, the majority of potentiated synapses revert to a depotentiated state within few hours (*1, 4*). This two-step process, mediated by the immediate early gene (IEG) Arc, results in harmonized bidirectional changes affecting multiple neighboring synapses, potentially underlying co-operative regulation of neuron-level plasticity (*4*). We hypothesized that shared local mechanisms may extend to coordination of synaptic plasticity over multiple dendrites and may be mediated by extracellular matrix (ECM) microenvironments.

To test this hypothesis, we focused on ‘CS6 Clusters’ (CS6Cs), large glia-derived proteoglycan aggregates with a predominant chondroitin sulfation (CS) in position 6 of N-acetylgalactosamine (CS6 moiety) (*6-8*). CS6Cs present as round structures, 50-200 μm in diameter, surrounding several dendrites and densely populating cortical and subcortical brain regions in the rodent, non-human primate and human brain (Figs.1A-C, 2 B, E) (*6-9*). Support for their role in synaptic plasticity comes from growing evidence for proteoglycans, and associated ECM molecules, as key functional components of synaptic extracellular scaffolding, linking pre- and post-synaptic terminals and powerfully regulating the actin cytoskeleton and synaptic structural plasticity (*10-15*). Specifically, CS6 positively modulates synaptogenesis, enhancing establishment of synaptic contacts (*16-19*). Based on this evidence, we hypothesized that CS6Cs represent segregated microenvironments encompassing several adjacent dendrites, potentially contributing to the coordination of plasticity in multiple synapses.

In primates and rodents, CS6Cs are detectable using antibodies raised against CS6 moieties (e.g. CS56 (*18*), used for this study). CS56 immunolabeling in the rodent, non-human primate and human brain shows that CS6Cs are morphologically conserved across species (Fig.1A-D). In all brain regions examined, CS6C morphology varies from a cluster of sharply-labeled converging processes (Rosette-CS6Cs; R-CS6Cs. Figs. 1A-C, 2B, S4A) to a cluster of diffuse puncta (Diffuse-C6Cs, D-CS6Cs; Figs. 1A-C, 2E, S4B). Operationally, we adopted the categorical definition of D-CS6C and R-CS6C (see Methods). However, these morphological types represent extremes of a spectrum, suggesting that D-CS6Cs and R-CS6Cs may represent different stages of the same structure.

**Fig. 1.**
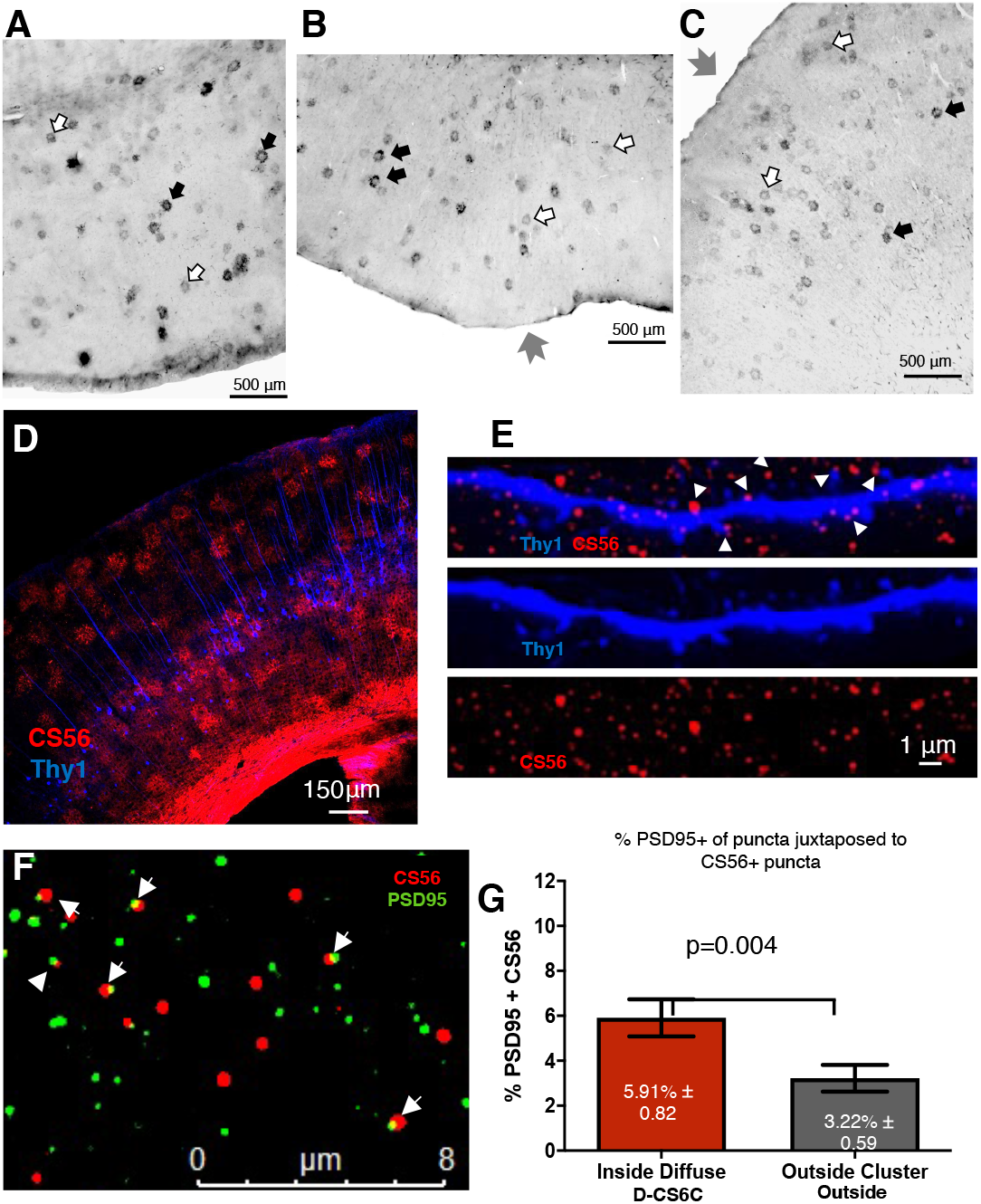
CS6Cs form microenvironments with dense CS6 accumulation within multiple glial processes and synaptic clefts. **(A)** CS6Cs (R-CS6Cs, black arrows; D-CS5Cs, white arrows) in the human entorhinal cortex, (**B**) non-human primate entorhinal and (**C**) mouse somatosensory cortices (grey arrows point to the pial surface). (**D**) CS6Cs and Thy1-positive dendrites (blue) in the mouse BCx, predominant in layers II-III and V. (**E**) CS6/CS56 punctae apposed to Thy-1-positive dendritic spines (arrowheads). (**F**) Juxtaposition (arrowheads) of PSD95-IR and CS56-IR puncta. (**G**) Percentage of PSD95-IR puncta juxtaposed to CS56-IR puncta within D-CS6C and outside adjacent areas.

**Figure 2.**
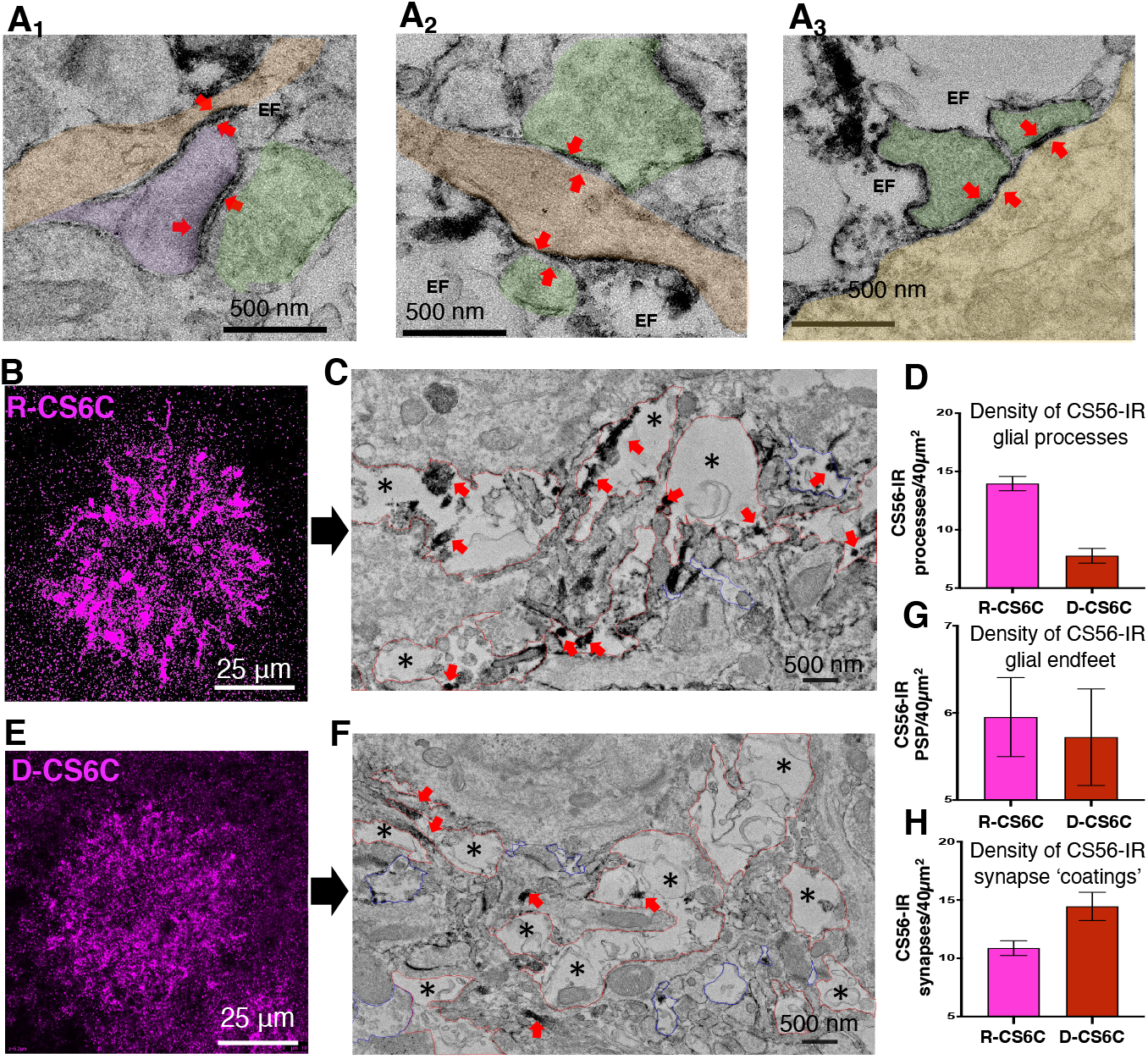
CS6Cs form microenvironments with dense CS6 accumulation within multiple glial processes and synaptic clefts. **(A**_**1-3**_) Examples of CS56-IR associated with synapses (red arrows; presynaptic terminals in green): **(A**_**1**_) on a dendritic spine (purple), **(A**_**2**_) a dendritic shaft (beige), and **(A**_**3**_) neuronal soma (yellow). **(B)** Examples of R-CS6C and (E) D-CS6C in the mouse BCx. **(C, F)** Distinct ultrastructural features of R- and D-CS6Cs (red arrow, CS56-IR; * glial processes). **(D)** R-CS6Cs present with predominant CS56-IR within glial processes. (**G**) R-CS6Cs and D-CS6C have similar densities of glial endfeet. **(F, H)** D-CS6Cs show higher densities of CS56-IR synapse ‘coatings’. Abbreviations: BCx, barrel cortex; EF, glial endfeet; IR, immunoreactive

In a first study, CS56 immunoprecipitation combined with proteomics on healthy human amygdala tissue was used to assess the CS6Cs proteoglycan composition and their potential binding partners. Our results show that the chondroitin sulfate proteoglycan (CSPG) versican is highly enriched in CS6Cs (Figs. S1-3, Table S1). CSPGs brevican and neurocan, as well as tenascin R and link proteins HAPLN1 and HAPLN2 are also represented, suggesting that CS6Cs are organized, complex ECM structures (Table S1). Notably, neurofilament light polypeptide was also detected in the CS56 immunoprecipitate (Table S1), consistent with interactions with extra- and intra-cellular synaptic elements.

To test the hypothesis that CS6Cs may contribute to local coordination of multi-synaptic plasticity, we focused on the mouse barrel cortex (BCx), a primary sensory region enriched in CS6Cs (Fig.1D). To test the association of CS6Cs with synapses, we used Thy1-eYFP mice, which show yellow fluorescent protein (YFP) fluorescence labeling in a subpopulation of pyramidal neurons, to visualize dendrites and dendritic spines (*20*). We found that CS6Cs encompass multiple apical dendrites in the BCx (Fig.1D) and that several Thy-1-labelled dendritic spines are juxtaposed to CS56-immunoreactive (CS56-IR) puncta (Fig. 1E). To assess the association between CS56-IR and excitatory synapses, we quantified PSD95-IR postsynaptic elements juxtaposed to CS56-IR puncta within and outside CS6Cs (Fig.1F,G). Within CS6Cs, 6% of PSD95-IR elements were contiguous to CS56-IR puncta. This percentage was significantly lower (3%) in an equivalent, same-layer area adjacent to CS6Cs (Fig.1G; Table S2). These findings indicate that CS56-IR elements are associated with synapses within CS6Cs, and to a lesser extent outside CS6Cs, where they are sparser.

We then assessed the ultrastructural relationship between CS6Cs and synapses, using transmission electron microscopy combined with pre-embedding CS56 immunolabeling. Within CS6Cs, CS56-IR product coats dendritic spines and fills the synaptic cleft (Figs.2 and S6,7). Axo-dendritic and axo-somatic synapses also showed CS56-IR coatings (Fig.2). CS56-IR within CS6Cs was also enriched within glial endfeet and glial processes, in association with intermediate filaments (Fig.2C,D). No CS56-IR was detected within neuronal cell bodies or dendrites (Fig.S6). Notably, key similarities and differences were detected between D-CS6Cs and R-CS6Cs (Fig. 2B-H). The number of CS56-IR glial endfeet is comparable in the two CS6C types. In contrast, R-CS6Cs contain higher numbers of CS56-IR glial processes. Conversely, D-CS6Cs contain higher number of synapses associated with CS6 coating. These results illuminate the CS6C structure, composed by glial processes and end-feet and extracellular coatings of CS6 moiety within the synaptic cleft. The differential distribution of CS56-IR may account for the distinct morphology of CS6Cs, i.e. R-CS6Cs characterized by a predominance of large glial processes, and D-CS6Cs characterized by predominant synaptic coatings and sparse or no glial processes, resulting in a punctate appearance (Figs.2, S4).

The association of CS56-IR with synapses is consistent with a well-established role of CSPGs in regulating synaptic plasticity (*10-15*). To test the hypothesis that CS6Cs may be involved in active phases of synaptic remodeling, we assessed whether sensory (whisker) manipulation affects CS6Cs within the mouse BCx. First, we compared the number of CS6Cs between hemispheres 1 week following unilateral whisker trimming (Fig.3A). The numerical densities (NDs) of CS6Cs was reduced in BCx layers II/III and V contralateral to whisker trimming as compared to the ipsilateral; no significant changes were observed in a control group (Figs.3B, S8). These results indicate that neuronal activation may be necessary for CS6C formation. Second, we tested whether sensory stimulation induces CS6C formation in the BCx, in parallel with the temporal progression of activity-dependent synaptic modifications shown to occur during the first 2 hours following a triggering stimulus (*3, 4*). Passive multi-whisker stimulation in anesthetized mice, shown to induce layer V-specific synaptic activation (*21*), was carried out by stimulating the left vibrissae (Fig.3D). Animals were sacrificed at 1 and 2 hours following the end of the stimulation. In the 1-hour group, R-CS6Cs, but not D-CS6Cs, were significantly increased in BCx layer V (Fig.3E-F). Conversely, D-CS6Cs, but not R-CS6Cs, were significantly increased in the 2 hours group (Fig.3 E,F). No significant difference was observed in two regions putatively unaffected by the stimulation, excluding the possibility of inter-hemispheric biases (Fig.S9). Together, these findings confirm both the necessity and sufficiency of neuronal activity for CS6C formation in cortex.

**Fig. 3.**
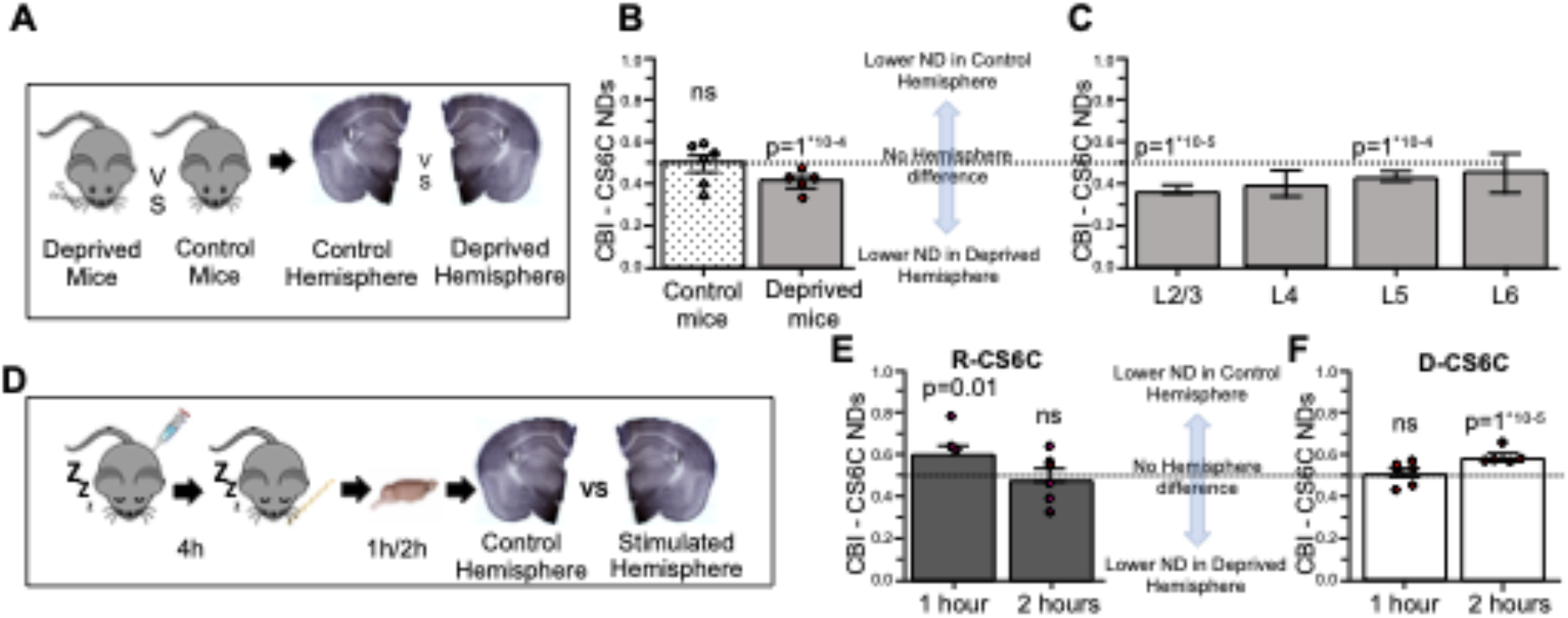
Sensory experience affects numerical densities (NDs) of CS6Cs in the Mouse BCx. **A)** Experimental design for sensory deprivation. **(B**,**C)** Sensory deprivation shifts CS6Cs NDs in favor of the non-deprived hemisphere in deprived mice, while controls show comparable number of CS6Cs across hemispheres. This result is driven by significant changes affecting predominantly layers 2/3 and 5. **(D)** Experimental design for sensory stimulation. **(E)** R-CS6Cs increase 1hr post-stimulation. **(F)** Conversely, D-CS6Cs selectively increase at 2h post-stimulation.

Ultrastructural and sensory stimulation findings indicate that R-CS6Cs and D-CS6Cs may represent temporally-related instances of the same transient structure. R-CS6Cs may arise soon after stimulation, as CS6 moieties are being transported by glial processes converging toward a group of neighboring synapses. R-CS6Cs morph into D-CS6Cs when CS6 moieties are secreted to form synaptic CS6 coatings. Notably, the temporal pattern of sensory stimulation-induced R-CS6Cs and D-CS6Cs increases parallels reports of local coordination of synaptic remodeling within 90 min from sensory stimulation, with synaptic structural long-term potentiation (LTP) plateauing at 2 hr, followed by structural long-term depression (LTD) (*4*). This timeline coincides with the formation of CS6 synaptic coatings on multiple dendrites (D-CS6C) during the structural LTD phase. To test this hypothesis, we used dendritic spine morphology as a structural proxy for synaptic LTP and LTD, corresponding to larger and narrower spine heads, respectively (*4, 22, 23*). In Thy1-eYFP mice, ‘mushroom’ spine width and length measures were compared within R-CS6C and D-CS6C and against control spines located outside the CS6Cs.

Our results show that, within CS6Cs, the correlation between length and width of spine heads is stronger with respect to control spines. Notably, CS6Cs do not contain the subpopulation of ‘narrow-head’ spines detected outside CS6Cs (Fig.4). Spines within R-CS6Cs have significantly larger heads and decreased head length with respect to control spines. The length-to-width ratio was ∼1 (Fig.4A), while control spine heads are significantly more elongated, corresponding to a ratio of ∼1.4. Conversely, spines within D-CS6Cs showed a significant decrease of head width and length as well as neck length (Fig.4B). The spine head length-to-width ratio within D-CS6Cs was ∼1.1, significantly different from both R-CS6Cs and control spines (Fig.4C,D). These results show that dendritic spine heads are significantly enlarged, putatively potentiated, within R-CS6Cs while their size is reduced within D-CS6Cs, putatively corresponding to the LTD phase. These findings support the hypothesis that R-CS6Cs and D-CS6Cs represent segregated active sites of synapse structural remodeling, potentially corresponding to LTP and LTD phases, respectively (*24*). The formation of CS6 synaptic coatings within D-CS6Cs corresponds, and may contribute, to coordinated synaptic de-potentiation of multiple synapses.

**Fig. 4.**
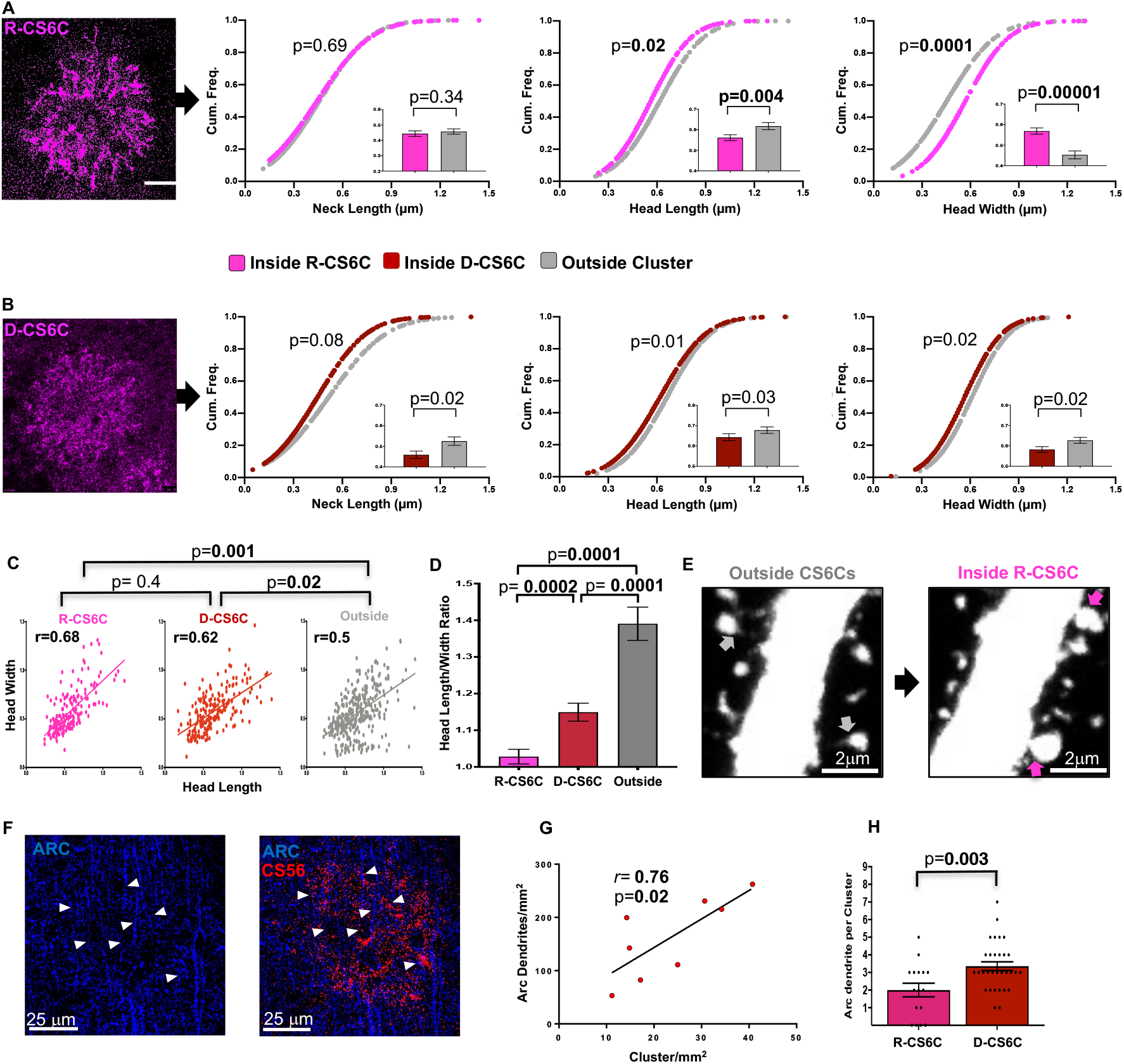
CS6Cs present subcellular and molecular features associated with activity-dependent plasticity. A) Within R-CS6Cs, dendritic spines present wider and shorter heads, but unaltered necks compared to outside CS6Cs. B) Within D-CS6Cs, dendritic spines are smaller compared to spines outside CS6Cs. C) Correlation between head length and width is stronger within CS6Cs. D) Spines within R-CS6Cs are stubbier than outside CS6Cs while spines within D-CS6Cs present with intermediate geometrical relationship. E) Confocal photomicrographs showing spines with large heads and short necks within a R-CS6C (magenta arrows) - outside CS6C spines have narrower heads and longer necks (gray arrows). F) Confocal photomicrographs showing multiple ARC-IR dendrites (white arrowhead) within a D-CS6C in the mouse BCx. G) Number of CS6Cs are positively correlated with ARC-IR dendrites within the BCx. H) D-CS6C contain higher numbers of ARC-IR dendrites compared to R-CS6.

To provide further support for the above hypothesis, we assessed dendrite expression of ARC, an IEG required for spine structural LTD and to coordinate multisynaptic plastic changes (*4*). Dual immunofluorescence labeling for ARC and CS56 (Fig. 4F) in a cohort of naïve mice showed a strong positive correlation between number of CS6Cs and ARC-positive dendrites within the BCx (Fig.4G). The number of ARC-IR dendrites within D-CS6Cs was significantly higher with respect to R-CS6Cs, indicating an increase of dendritic ARC expression within D-CS6Cs (Fig.4H). These findings further corroborate the idea that D-CS6Cs may be associated with a phase of structural synaptic depression.

Taken together, these studies provide the first evidence that CS6Cs are involved in spatially-coordinated, stimulus-driven structural plasticity. They support the hypothesis that CS6Cs are transient structures involved in coordinated plasticity of multiple synapses within a segregated space (Fig. S11). This mechanism may represent a novel modality of interaction between glial cells, ECM and synaptic elements. We propose that stimulus-induced glia-derived CS6 moieties may become dynamically and transiently incorporated into the extracellular synaptic scaffolding, potentially contributing to structural synaptic changes. These results introduce a new player into the field of neuroplasticity, a novel ECM structure potentially contributing to locally-coordinated synaptic plasticity. Abundant presence of CS6Cs in the human brain and decreases in schizophrenia and bipolar disorder (*9*) support a role for these structures in adult human neural plasticity and pathophysiology of brain disorders.

## Funding

**SB:** NIH R01MH104488, NIH R01MH120991; **GC**: Rappaport Mental Health Research Fellowship 2018/19; **TK**: NIH R21HD088931, R21HD097524, R21MH117609 and R01HD102974; **RM:** NIMH R01 MH107487 and MH121102 and **KJR**: NIH R01 MH108665 and MH120991, NIH P50 MH115874.

## Author contributions

GC carried out most of the research project and contributed to the conceptualization of this project and writing of manuscript, PD contributed to data collection and analyses, S O’D and RM carried out proteomics studies in human and contributed to the conceptualization of this work, TK contributed to the conceptualization, experimental design and manuscript preparation, LB and YB contributed to experiments on sensory stimulation and manuscript preparation; CB carried out the electron microscopy work, Anne Boyer-Boiteau contributed to the research project, Kerry J. Ressler contributed to the conceptualization of this project, resources and manuscript preparation, SB contributed to the conceptualization of the project and its supervision, data analysis, resources and manuscript preparation.

## Competing interests

TK has received consulting income from Alkermes Inc, Waltham, MA unrelated to this work. Authors declare no competing interests.

## Data and materials availability

All data is available in the main text or the supplementary materials.

## Supplementary Materials

Materials and Methods

Figures S1-S11

Tables S1-S6

References (*25-33*)

## Notes

### Competing Interest Statement

The authors have declared no competing interest.

