## Supplementary Methods and Data for "Proteoglycan Clusters as a Site of Coordinated, Multi-Dendritic Plasticity"

### SUPPLEMENTAL MATERIAL

### MATERIAL AND METHODS

#### *Human Postmortem Studies*

CS56-Immunoprecipitation and Mass Spectrometry Analysis

• *Tissue preparation*

Fresh-frozen human postmortem amygdala tissue samples from healthy donors were obtained from the Harvard Brain Tissue Resource Center (HBTRC; McLean Hospital, Belmont MA). Several regions from each brain were examined by a neuropathologist. The cohort used for this study did not include subjects with evidence for gross and/or macroscopic brain changes, or clinical history, consistent with cerebrovascular accident or other neurological disorders. Subjects with Braak stages III or higher were not included. None of the subjects had significant history of substance dependence within 10 or more years from death, as further corroborated by negative toxicology reports. Absence of recent substance abuse is typical for samples from the HBTRC, which receives exclusively community-based tissue donations. Frozen tissue blocks containing the amygdala were placed in a cryostat maintaining a constant temperature between -13$^{o}$ and -15 $^{o}$C. The borders of the amygdala were lightly 'etched' using a surgical knife so that collected sections did not contain surrounding brain regions.

Sections were homogenized in buffer (320 mM sucrose, 5 mM Tris, pH 7.0) containing HALT protease inhibitor cocktail (Thermo Fisher Scientific, Carlsbad, CA). For affinity purification, sample was incubated overnight at 4°C with 1% TritonX-100 and 0.5% SDS. Samples for immunoblotting were boiled for 10 mins at 70°C in sample buffer (6X solution: 4.5% SDS, 15% β-mercaptoethanol, 0.018% bromophenol blue and 36% glycerol in 170 mM Tris-HCl, pH 6.8). Sample protein concentration was determined by a BCA protein assay (Thermo Fisher Scientific, Waltham, MA).

• *Immunoblotting*

Twenty micrograms of protein were loaded onto 4-12% Novex NuPage Bis-Tris 10-well gels (1.5mm) (Life Technologies, Waltham, MA). Gels were run at 180 V for 1 hour and then transferred to PVDF membranes by Bio-Rad semidry transblotters (Bio-Rad, Hercules, CA, USA) as described previously (Shan et al, 2014). All membranes were blocked by LiCor blocking buffer (LiCor, Lincoln, NE, USA) for 1 hour at room temperature. Membranes were probed overnight at 4°C for all primary antibodies. The antibodies used were: 1:500 mouse monoclonal CS-56 (ab11570, Abcam, Cambridge, MA), 1:200 rabbit polyclonal Versican (sc-25831, Santa Cruz Biotechnology, CA, USA), 1:200 rabbit polyclonal (ab36861, Abcam, Cambridge, MA, USA). Following 3 x 10 minute washes in 1X PBS, membranes were incubated with appropriate IR-Dye 670 or 800cw labelled secondary antibodies (1:10,000) in LiCor blocking buffer for 1 hour at room temperature. Following 3 x 10 minute PBS washes, membranes were scanned using a LiCor Odyssey scanner.

• *CS56 affinity purification*

CS56 affinity purification (AP) was performed by immunoprecipitation, using the Novex Life Technologies Dynabeads kit according to manufacturer's guidelines, with some modifications. Briefly, 5 µg of antibody were conjugated per 1 mg of M-270 Epoxy beads (Life Technologies, Oslo, Norway) overnight at 37°C on a shaker. Beads were conjugated with CS-56 (ab11570, Abcam Cambridge, MA, USA) or the corresponding isotype control IgM (sc-3381, Santa Cruz Biotechnology, Santa Cruz, CA, USA). The beads were rinsed 4 x 10 minute in 1X PBS-T. Ten milligrams of antibody-conjugated beads were incubated overnight at 4°C with 742 µg of tissue homogenate. The following day, the supernatant was collected and retained for immunoblotting.

Beads were rinsed (4 x 10 minute washes in 1X PBS-T) and the bound proteins were eluted in 30 µl of elution buffer (1 N ammonium hydroxide, 0.5 M EDTA, pH 11.6).

• *Mass spectrometry*
CS56 affinity purified elutions were boiled at 70°C for 10 minutes in sample buffer. The sample was loaded on a 1.5 mm, 4-12% gradient gels and electrophoresed until the sample ran 1.5 cm into the gel. Molecular weight markers (Thermo Spectra 26623) were run between the samples to indicate the protein-containing region of the gel. The gel was fixed in 50% ethanol/10% acetic acid overnight at RT, then washed in 30% ethanol for 10 min followed by two 10 min washes in MilliQ water (MilliQ Gradient system) and finally scanned on an Epson V700 scanner to record an image of the gel. The lanes were cut out of the gel, cut into small (~2mm) squares and were subjected to in-gel tryptic digestion and subsequent recovery of peptides.
Nano liquid chromatography coupled to electrospray tandem mass spectrometry (nLC-ESI-MS/MS) analyses were performed on a 5600+ QTOF mass spectrometer (Sciex, Toronto, On, Canada) interfaced to an Eksigent (Dublin, CA) nanoLC. ultra nanoflow system. Each acquisition method was analyzed in technical triplicate. Protein was loaded (via an Eksigent nanoLC.as-2 autosampler) onto an IntegraFrit Trap Column (outer diameter of 360 µm, inner diameter of 100, and 25 µm packed bed) from New Objective, Inc. (Woburn, MA) at 2 µl/min in formic acid/$H_2O$ 0.1/99.9 (v/v) for 15 min to desalt and concentrate the samples. For the chromatographic separation of peptides, the trap-column was switched to align with the analytical column, Acclaim PepMap100 (inner diameter of 75 µm, length of 15 cm, C18 particle sizes of 3 µm and pore sizes of 100 Å) from Dionex-Thermo Fisher Scientific (Sunnyvale, CA). The peptides were eluted using a variable mobile phase (MP) gradient from 95% phase A (Formic acid/H2O 0.1/99.9, v/v) to 40% phase B (Formic Acid/Acetonitrile 0.1/99.9, v/v) for 35 min, from 40% phase B to 85% phase B for 5 min and then keeping the same mobile phase composition for 5 additional min at 300 nL/min. The nLC effluent was ionized and sprayed into the mass spectrometer using NANOSpray® III Source (Sciex). Ion source gas 1 (GS1), ion source gas 2 (GS2) and curtain gas (CUR) were respectively kept at 12, 0 and 35 vendor specified arbitrary units. The mass spectrometer method was operated in positive ion mode and the interface heater temperature and ion spray voltage were kept at 125ºC, and at 2.6 kV respectively. The data was recorded using Analyst-TF (version 1.7) software.

• *Data analysis*
Protalizer software (Vulcan Analytical) was used to analyze mass spectrometry data as previously described (*1*). Briefly, peptide and protein identifications were made using the X! Tandem Sledgehammer search engine (version 2013.09.01.1) with a 50 ppm precursor and fragment-ion mass tolerance. Potential modifications included in the searches were phosphorylation at S, T, and Y residues, N-terminal acetylation, N-terminal loss of ammonia at C residues, and pyroglutamic acid at N-terminal E and Q residues. Carbamidomethylation of C residues was searched as a fixed modification. A maximum of two trypsin miscleavages were included in the analysis and protein and peptide maximum valid expectation scores were set to 0.005. Only 'top hit' peptides were further analyzed by the pipeline. Redundant peptide identifications with the same charge and modification state were eliminated for downstream analysis in each file except for one scan corresponding to the largest sum fragment-ion intensity according to X! Tandem.

***Experimental animal studies***
All experimental mice were group housed (3–5 mice per cage) on a 12:12hr light-dark cycle in a temperature-controlled colony room and had unrestricted access to food and water. All procedures were conducted in accordance with policy guidelines set by the National Institutes of Health and were approved by the McLean Institutional Animal Care and Use Committee, the Animal Welfare Committee of the University of Trento, and the Italian Ministry of Health.
<u>Sensory deprivation, electron microscopy, and ARC studies</u>. Young adult (Postnatal day [P] 70)

C57-black wild type male mice were purchased from The Jackson Laboratory. Mice were let to habituate to the housing conditions in the McLean Hospital animal facility for a week before being used for experiments.
Sensory stimulation study. C57BL/6 mice were originally purchased from Charles River (Italy) and bred in the animal facility of the University of Trento (Italy). Adult C57BL/6 male mice (P 70-95; weight 23-28 g) were included in the study.

Dendritic spine analysis. Eight naïve Thy1-eYFP male mice (The Jackson laboratory), age P70-90 were used in the study.

Perfusion and tissue processing

• *Electron Microscopy Studies.*

Mice were deeply anesthetized with isoflurane and perfused transcardially with 50 ml phosphate buffer/saline (PBS) followed by 50ml PFA-glutaraldehyde-NaCacodylate (4% PFA, 0.1% glutaraldehyde, 0.1M Sodium Cacodylate, 0.21mg/ml $CaCl_2$ 0.1M Sucrose). Brains were postfixed in PFA-NaCacodylate (4% PFA, 0.1M Na Cacodylate, 0.21mg/ml $CaCl_2$, 0.1M Sucrose) for 2 hours. Fixed brains were kept overnight at 4°C in PBS and sectioned on a vibratome (Leica VT1200S) at 40µm thickness. Sections for each animal were collected in PBS and processed 24h later for immunohistochemistry.

• *Light and Confocal Microscopy Studies.*

Mice were deeply anesthetized with isoflurane (urethane for sensory stimulation experiment, see below) and perfused intracardially with a 50ml 0.1M phosphate buffer (PB), followed by 50 ml of 4% paraformaldehyde (PFA; pH 7.4) (for sensory deprivation experiments 100ml of 4% PFA was used). Brains were harvested and post-fixed overnight in 10ml of 4% PFA, then switched to a cryoprotectant solution (80% PB, 20% glycerol with 0.1% sodium azide) and stored at 4°C in cryoprotectant solution. Cryoprotected brains were sectioned on a freezing sliding microtome (American Optical 860, Buffalo, NY, USA) and serial sections (30µm thickness) were collected in 24 separate compartments (each one containing a complete antero-posterior representation of the whole brain) and stored at 4°C in cryoprotectant solution.

Immunohistochemistry

• *Single Immunolabeling for Light Microscopy*

Step 1. Free-floating tissue sections were washed for 90 mi to rinse out the cryoprotectant, then pre-treated with antigen retrieval solution overnight at 37°C (1.8mM DTT, 1.8mM EDTA, 12mM Citric Acid, 43mM, 43mM $Na_2HPO_4$, 0,05% bovine serum albumin (BSA) in deionized-distilled $H_2O$). Sections were then incubated in 0.03 % $H_2O_2$ solution for 30 min, followed by incubation in blocking solution (2% BSA (fraction V)) for 90 min. Tissue was then transferred to primary antibody solution (2% BSA solution; see primary antibody concentrations below) and incubated at 4°C for 60 hr.
Step 2. Sections were incubated with a biotinylated secondary antibody (1:500) of for 2 hr, followed by streptavidin conjugated with horse-radish peroxidase (Zymed, San Francisco, CA, USA. 1:5000) for 2 hr. Sections were then rinsed in PB and incubated with nickel-enhanced diaminobenzidine/peroxidase (0.02% diamino-benzidine (*Sigma,* D5905-100TAB*)*, 0.08% nickel-sulfate, 0.007% $H_2O_2$ in PB). The colorimetric reaction was visually monitored and stopped by washes in PB. Tissue sections were mounted on gelatin-coated slides, dried overnight, dehydrated through serial ethanol-xylene steps and coverslipped with a xylene-based mounting media (Thermofisher Cytoseal, 8310-16). All solutions were made in 0.1 PBS containing 0.2% Triton X-100 (Fisher, AC215680010) (PBS-Tx) and all steps were preceded and followed by washes consisting of 3 consecutives 5 min rinses in PBS-Tx, unless otherwise specified.

• *Single and Double Immunofluorescence Labeling*
Step 1. As described above
Step 2. Tissue sections previously incubated in the primary antibody solution are rinsed, then placed in a fluorophore-conjugated secondary antibody solution (1:300) for 4 hr. Sections are then washed 5 min. in PB, mounted on gelatin coated slides, dried for 1 hr, coverslipped with fluorescent mounting medium (Southern biotech 0100-01) and stored in the dark at 4°C. All solutions were made in 0.1 PBS containing 0.2% Triton X-100 (Fisher, AC215680010) (PBS-Tx) and all steps were preceded and followed by washes consisting of 3 consecutives 5 min rinses in PBS-Tx, unless otherwise specified.

• *Pre-embedding Immunolabeling for Electron Microscopy*
Free-floating tissue sections were rinsed in PB, treated with 0.3% Hydrogen peroxide (*Fisher, H325100)* for 15 min, and then incubated in a solution of 0.05% sodium borohydride *(Fisher, AC448501000),* 0.1% Glycine *(Sigma, G-7403)* for 30 min. Next, the sections were incubated in blocking solution (4% Bovine BSA *(Fisher, BP1605100),* 10% Goat serum *(Gibco, 16210-064),* 0.01% Triton X-100 *(Fisher, AC215680010)*) for 90 min, then transferred in primary antibody solution (1:1000 CS56, 2% BSA, 2% Goat serum, 0.01% Triton X-100) for 2 days. A subset of sections was incubated without primary antibody as a negative control. The sections were then incubated in biotinylated anti-mouse IgM secondary antibody (*Vector labs, BA-2020)* 1 to 500µl) for 2 hr, followed by incubation in Streptavidin-HRP (*Invitrogen, 434323)* (1:5000l) for another 2 hr. Finally, tissue sections were washed with PB and incubated with 3,3'-Diaminobenzidine Tetrahydrochloride *(Sigma,* D5905-100TAB*)* 0.5mg/ml in 0.1M Sodium Cacodylate, 0.005% hydrogen peroxide for 20 min. The reaction was stopped and sections washed with PB and processed for EM (see below). All solutions were made in 0.1 PBS and all steps were preceded and followed by washes consisting of 3 consecutives 5 min rinses in PBS, unless otherwise specified.

• *Primary Antibodies*
***CS56.*** The antibody used is a mouse monoclonal IgM (Sigma-Aldrich, C8035) made using vertebral membranes from chicken gizzard fibroblasts as an immunogen. The CS structure immuno-reactive for CS56 has been identified as an A–D sequence in reducing octosaccharide units on both CS-C (CS-6) and CS-D (CS-2,6) chondroitin sulfate[25,26]. CS56 antibody was previously shown to detect CS6 clusters in human and mouse tissue[8-10]. The antibody was diluted 1:6000µl for light microscopy and 1:3000µl for immunofluorescence.
***Arc/Arg3.1.*** The antibody used is an affinity-purified rabbit polyclonal IgG antibody (Proteintech, 16290-1-AP). This antibody was previously used to detect ARC protein localization in dendritic arbors[27].The antibody was diluted at 1:1000µl for immunofluorescence.
***PSD95.*** The antibody used is a polyclonal IgG antibody made in goat (Abcam, AB90426). Antibody was diluted 1:500µl for immunofluorescence.

• *Electron Microscopy*
Immunostained tissue sections were postfixed in 1% OsO4, dehydrated in graded ethanol series and extra-dry acetone, and flat embedded in Embed 812 resin. Ultrathin sections (~80 nm) were collected on Formvar-coated slot grids. A set of sections was left unstained and a second set was post-stained with uranyl acetate (saturated solution) and Reynold's lead citrate, before being analyzed. Sections were imaged on a JEOL JEM-1200 EX II with an 1k CCD camera, and on a Tecnai F20 (200 keV) transmission electron microscope (FEI, Hillsboro, OR); the images were collected using a 2K × 2K charged-coupled device (CCD) camera, at 19,000X magnification (1.12

nm pixel size). Whole CS6Cs photocomposites we acquired as automated montages of overlapping high-magnification images using the microscope control software SerialEM[28].

Behavioral Paradigms

• *Sensory Deprivation (whisker trimming)*

C57-black wild type male mice were assigned to a control group (n=6) and a sensory-deprived group (n=5). Sensory-deprivation was achieved by closely trimming the right facial vibrissae for 7 consecutive days every 24 hours under brief isoflurane anesthesia, according to published protocols[23]. Minimal whisker regrowth was observed within 24 hours following each trimming session. Control mice were briefly anesthetized daily to maintain similar experimental conditions. On day 8, animals were sacrificed and perfused as described above.

• *Sensory Stimulation (whisker stimulation)*

C57-black wild type male mice were assigned to two groups, one sacrificed at 1h (n=6) and one at 2 hours (n=6) following sensory stimulation. Unilateral whisker stimulation was performed using a protocol described in Chelini et. al, 2019[24]; the ipsilateral hemisphere served as a control. All mice were anesthetized by intraperitoneal injection of a 20% solution of urethane in sterile double distilled water (1.6 g/kg body weight), placed on a 37°C warming pad, and head-fixed on a stereotaxic frame. Urethane anesthesia was chosen as it preserves whisker-dependent activity in the somatosensory cortex[4]. Mice were kept under anesthesia for 4 hours prior to whisker stimulation. Immediately before whisker stimulation, a subcutaneous saline injection (100-200 μl) was administered to maintain hydration. Additional anesthetic at 10% of the initial dose were administered as needed. Whisker stimulation was performed in three consecutives 5 minute sessions with 1 minute intervals. Each session consisted of unilateral stimulation on the left side by continuous whiskers deflection using a wooden stick. Mice were kept under constant anesthesia and sacrificed at two different time points, 1hr and 2hr, following the end of whisker stimulation.

Data Collection and Analysis

Data collection for all studies was carried out by investigators blind to experimental conditions. Slides and images were coded by an investigator not involved in data collection. All statistics analyses were carried out using Prism8 software (GraphPad Software San Diego, CA)

• *CS6C Classification*

CS6Cs present with distinct morphological characteristics, such a round shape with a diameter of 70-100μm (in rodents) often enveloping several unstained cell bodies (examples in Fig. S4, S5). CS6C morphological features, such as sharply labeled processes and diffuse punctate labeling, are present to different degrees across a spectrum. For practical quantification purposes, we assigned CS6Cs to two distinct categories, i.e. R-CS6C and D-CS6C, depending on their predominant morphological features. CS6Cs with negligible punctate labeling and predominant sharply labeled converging processes forming a Rosette were classified as R-CS6C (Fig. 2B, S4A). Conversely, CS6Cs with a predominant pattern of diffuse immunolabeled puncta was categorized as D-CS6C (Fig. 2E, S4B). Note that this latter category also included intermediate morphologies, i.e. CS6Cs containing some immunolabeled processes intermingled with dense punctate labeling. EM images are consistent with this classification, showing that peri-synaptic coatings and glial endfeet represent the majority of immunolabeled elements in this intermediate CS6C types.

• *Stereology-Based CS6C Quantification for Sensory Studies*

Zeiss Axioskop 2 Plus interfaced with StereoInvestigator 6.0 (Micro-Brightfield, Willinston, VT, USA) was used for CS6C quantification. Using StereoInvestigator software, the borders of the

BCx were traced according to Allen Brain Atlas (https://mouse.brain-map.org/static/atlas), and layers II-III, IV, V and VI according to cytoarchitectonic characteristics. As control region for sensory stimulation experiments, layer V of the Motor Cortex was also included in these analyses. All serial sections within a compartment, representing the full rostro-caudal extent of the BCx (section interval 720μm), were included in the analyses, thus respecting the equal opportunity rule. Within each layer, all morphologically identifiable CS6Cs were classified as R- or D- CS6Cs on the basis of their morphological features and counted. Numerical densities (NDs) were calculated as $ND = \sum N / \sum V$, where N is the sum of elements within a region of interest, and V is the total volume (in $mm^3$) of the region of interest (estimated over the sectioning interval). Total number (TN) of IR elements was calculated as $N = i \times \Sigma n$ where $\Sigma n$=sum of the cells counted in each layer/mouse, and i is the section interval (24, i.e. the number of serial sections between each section and the next within each compartment), as described previously in detail[29].

Two measures were used to compare inter-hemispheric differences:

- Inter-hemispheric differences in the NDs of CS6Cs were compared using a 2-tailed paired t-test.
- Contralateral bias index (CBI) was calculated for each animal using the formula: ND Left Hemisphere/(ND Right Hemisphere + ND Left Hemisphere) and compared to the predicted average (CBI= 0.5) using a Z-test. Values significantly higher than 0.5 reflect higher CS6S ND in the left hemisphere. Values significantly lower than 0.5 reflect higher CS6C ND in the right hemisphere. No statistical difference with the expected average (0.5) reflects similar CS6C NDs across hemispheres.

To rule out the possibility of intrinsic inter-hemispheric biases in the sensory stimulation experiment, resulting in increased clusters NDs in either the left or right hemisphere under no direct stimulation, we quantified inter-hemispheric differences in two regions putatively unaffected by sensory stimulation, i.e. Motor cortex (layer V) and BCx (layer VI). No significant changes of CS6C ND between hemispheres (data not shown), nor significant shift of CBI from the predicted average (Fig. S9) was detected.

• *Electron Microscopy Analysis*

One prototypical R-CS6C and one D-CS6C were acquired from an adult (P80) wild-type mouse and compared in this study. High magnification (19,000x) montages of whole CS6Cs in not-counterstained sections were deconstructed in multiple quadrants (8.6x4.5μm) and analyzed using StereoInvestigator 6.0 software. Inside each field, CS56 immunoreaction products were manually quantified. Glial processes were recognized according to Peters et al.[30] (Fig. S6, 7). Each independent immunoreaction product within a glial cell process was quantified as a single element (Fig. S6,7). When positive immunoreaction was observed within the edge of a glial cell process adjacent to a pre-or-post synaptic compartment the element was categorized as “glia end-foot” (Fig. 2 and S6. Blue stars).

Synapses, identified by their distinct pre- and post-synaptic elements, were counted as positive (CS6-coated) if they showed clearly distinguishable black immunoreaction product within the synaptic cleft and/or coating the pre- and post-synaptic elements (Fig. 2 and S6). These elements were quantified by counting each instance of distinct CS56 immunoreaction product within them in each quadrant (Fig. S6). Statistical differences between R-CS6Ce and D-CS6C with respect to numbers of CS56-IR elements (glial processes, glial end-feet and synapses) were compared using a t-test.

• *ARC-CS56 double labeling analyses.*

In the BCx of 4 mice, we counted ARC-IR dendrites within 17 R-CS6Cs and 33 D-CS6Cs. A Zeiss Axio Imager M2 with a Lumencor SOLA LED lamp interfaced with StereoInvestigator 10.0

(Microbrightfield Inc., Willinston, VT) was used for these analyses. As a first step, total number of CS6Cs and ARC-immunoreactive (IR) dendrites were quantified layer II/III of the BCx on dual antigene immunolabeled sections. Pearson correlation was used to infer correlations between numerical densities of CS6Cs and ARC-IR dendrites. ARC-IR dendrites were then counted within R-CS6C and D-CS6C. Statistical differences of ARC-IR dendrites contained within R-CS6C vs D-CS6C were assessed using a t-test.

• *Synapse Analyses*
A confocal laser scanning microscope Leica TCS SP8, equipped with a HC PL APO 100 x/1.40 oil objective interfaced with Leica LAS-X software was used to acquire image z stacks for all synapse-related studies below. Images were recorded at high resolution (1024 pixels square), 400 Hz scan speed, in sequential imaging mode. Excitation/emission wavelengths used were: 490/520 for Alexa 488 fluorophore, 640/670 for cy5 647 fluorophore.

• *CS56-PSD95 Co-Localization Analyses*
Z-stack (0.5μm optical thickness) images were acquired within D-CS6C clusters and size-matched areas outside a CS6C within the BCx at 100x magnification. The choice of quantifying D-CS6C was based on EM data showing predominant CS6 synaptic coats within D-CS6C as well as the fact that prominent glia processes immunolabeling within R-CS6C rendered synaptic quantification unreliable. Acquisition parameters were kept consistent across image acquisitions. Individual stacks were quantified using StereoInvestigator 6.0 software by an investigator blind to the experimental conditions. Quantification was performed using a standardized counting strategy: First, total number of individual CS56- and PSD95-IR punctate elements were quantified separately on single channels for each image stack. Second, dual channel images were used to quantify the number of appositions between PSD95- and CS56-IR puncta. Partially overlapping puncta labeling with less than 1-pixel distance between the PSD95- and CS56-IR puncta were included. Percentages of PSD95-IR puncta apposed to CS56-IR puncta inside and outside D-CS6Cs were compared using a t-test.

• *Dendritic Spine Morphological Analyses*
Coronal sections including the BCx were obtained from Thy1-YFP mice[18]. Z-stack images of YFP-labeled dendrites were acquired in layer II/III at 100x magnification. For each dendrite analyzed, two sets images were collected, one of a stretch located within a CS6C and a second of a stretch of same dendrite located outside the CS6C. We refer to spines located on dendrite stretches outside CS6Cs as 'control' spines. This strategy was designed to control for potential variability between dendrites within and outside CS6Cs. Furthermore, because dendritic spine structural dynamics depend in part on their proximity to the soma, we sampled dendrite stretches outside the proximal and distal side of CS6Cs with respect to the cell body. Images were analyzed offline using Leica LAS-X image analyzer software.

Before proceeding with mushroom spines morphometric analysis, spines were quantified and classified into five different categories according to their morphological features: mushroom, stubby, thin, filopodia and ramified as previously described by Hering and Shang, 2001[31]. Only a small fraction of ramified and filopodia were observed, so they were not included in this analysis. Dendritic spine density was defined as *number of dendritic spines/length of dendritic stretch (in μm)*. Dendritic segments located within a CS6C were compared with segments of the same dendrites residing outside using a 2-tailed paired t-test. No significant difference was observed in dendritic spine density for neither mushroom, stubby or thin spines (Fig. S10).

Dendritic spine morphological changes have been widely used as a structural proxy for synaptic potentiation and de-potentiation[21, 22]. Specifically, shorter and wider spines are interpreted as potentiated synapses, while longer and thinner spine may represent a de-potentiated state[21, 22].

For these studies, we measured the head length, head width and neck length of each mushroom spine, using the measuring tool item included in the Leica LAS-X software (Fig. S10). *Neck length* was defined by the length from the most medial point of the base of the neck to the most medial point of the most distal aspect of the neck. *Head length*, similarly, was measured from the most medial and proximal aspect of the head to the most medial and distal aspect. The *width of the head* was measured at the widest aspect of the head. Total spine length was obtained combining neck and head length. Kolmogorov-Smirnov test was used to assess differences in the frequency distribution between different spine populations. A Mann-Whitney test was used to assess average differences for each measurement. Linear correlation between spine head length and width were assessed using Pearson correlation and differences in the linear correlation were compared across groups using a Z-test. Ratio between spine head length and width was compared between R-CS6C, D-CS6C and control spines using a t-test.

### SUPPLEMENTAL SUPPORTING DATA

Supplemental Tables

| Accession no. | Protein name | Gene | # Peptides identified |
|---|---|---|---|
| P13611 | Versican core protein | VCAN | 147 |
| O14594 | Neurocan core protein | NCAN | 17 |
| Q96GW7 | Brevican core protein | BCAN | 3 |
| P10915 | Hyaluronan and proteoglycan link protein 1 | HAPLN1 | 8 |
| Q9GZV7 | Hyaluronan and proteoglycan link protein 2 | HAPLN2 | 6 |
| P07196 | Neurofilament light polypeptide | NEFL | 10 |
| Q86YZ3 | Hornerin | HRNR | 9 |
| P09543 | 2',3'-cyclic-nucleotide 3'-phosphodiesterase | CNP | 4 |
| P19823 | Inter-alpha-trypsin inhibitor heavy chain | ITIH2 | 3 |
| Q92752 | Tenascin-R | TENR | 3 |

**Table S1. Identification of CS-56 protein complex following affinity purification (AP) using mouse anti CS-56 antibody**. Proteins are identified by Swiss-Prot accession number, protein name and gene name. Ten proteins were identified as potential CS-56 interacting proteins. These proteins were found only in the CS-56 AP (proteins also identified in the IgM control AP were removed) and were identified with a minimum of 2 unique peptides.

| | Percentage of PSD95-IR elements juxtaposed to CS56-IR puncta | |
|---|---|---|
| | Mean | SEM |
| Outside CS56C | 3.22% | 0.59 |
| Inside CS56C | 5.91% | 0.82 |

**Table S2. Comparison of PSD95-IR elements juxtaposed to CS56-IR puncta within D-CS6Cs as compared outside CS6C**. Percentages of PSD95-IR puncta juxtaposed to CS56-IR puncta are significantly higher within CS6Cs as compared to an equivalent area adjacent to them (p=0.004). Measurements were carried on Z-stacks of images obtained by confocal microscopy on the BCx of naïve wild type mice (n=3). Five CS6Cs and 5 adjacent areas of the same size were counted for this analysis.

Table 3

| Group | Hemisphere/layer | Mean | SEM | CBI | CBI SEM | N mice | Paired t-test | Z-test (CBI) |
|---|---|---|---|---|---|---|---|---|
| **Control Mice** | Right Hem. | 4817.9 | 940.91 | 0.49 | 0.04 | 6 | 0.55 | 0.86 |
| | Left Hem. | 4281.81 | 378.37 | | | | | |
| | L2/3 Right Hem. | 1517.91 | 444.57 | 0.55 | 0.05 | 6 | 0.25 | 0.34 |
| | L2/3 Left Hem. | 1001.86 | 122.003 | | | | | |
| | L4 Right Hem. | 992.76 | 229.74 | 0.43 | 0.05 | 6 | 0.44 | 0.26 |
| | L4 Left Hem. | 1189.76 | 153.67 | | | | | |
| | L5 Right Hem. | 1223.18 | 168.76 | 0.51 | 0.03 | 6 | 0.4 | 0.71 |
| | L5 Left Hem. | 1081.78 | 83.98 | | | | | |
| | L6 Right Hem. | 1084.03 | 227.44 | 0.54 | 0.05 | 6 | 0.77 | 0.92 |
| | L6 Left Hem. | 1008.4 | 107.09 | | | | | |
| **Sensory Deprived Mice** | Right Hem. | 4827.9 | 519.43 | 0.4 | 0.02 | 5 | **0.01** | **0.0001** |
| | Left Hem. | 3412.66 | 547.69 | | | | | |
| | L2/3 Right Hem. | 1583.93 | 257.9 | 0.61 | 0.01 | 5 | **0.01** | **0.00001** |
| | L2/3 Left Hem. | 969.3 | 154.02 | | | | | |
| | L4 Right Hem. | 1109.42 | 65.58 | 0.59 | 0.06 | 5 | 0.33 | 0.12 |
| | L4 Left Hem. | 848.61 | 227.53 | | | | | |
| | L5 Right Hem. | 1519.48 | 202.71 | 0.55 | 0.02 | 5 | 0.07 | **0.01** |
| | L5 Left Hem. | 1162.97 | 98.99 | | | | | |
| | L6 Right Hem. | 944.5 | 287.62 | 0.54 | 0.09 | 5 | 0.46 | 0.63 |
| | L6 Left Hem. | 675.97 | 113.04 | | | | | |

**Table S3.** Unilateral whisker trimming (1 week) significantly decreases CS6Cs-NDs in the contralateral BCx. This effect is predominantly driven by changes in layer (L) 2/3 and L5. No inter-hemispheric difference is observed in a group of untreated animals (controls).

Table 4

| CX area /layer | Survival time | Cluster type | Hemisphere | Mean | SEM | CBI | CBI SEM | Paired t-test | Z-test (CBI) |
|---|---|---|---|---|---|---|---|---|---|
| **BCx L5** | **1 hr** | **R-CS6C** | Stimulated | 241.16 | 47.53 | 0.61 | 0.03 | 0.05 | **0.01** |
| | | | control | 160.6 | 39.66 | | | | |
| | | D-CS6C | Stimulated | 923.07 | 81.48 | 0.51 | 0.01 | 0.59 | 0.41 |
| | | | control | 813.74 | 68.83 | | | | |
| | **2 hr** | R-CS6C | Stimulated | 514.01 | 97.37 | 0.48 | 0.05 | 0.74 | 0.72 |
| | | | control | 545.06 | 85.43 | | | | |
| | | **D-CS6C** | Stimulated | 1418.57 | 152.33 | 0.58 | 0.01 | **0.002** | **0.00001** |
| | | | control | 1002.6 | 118 | | | | |
| **BCx L6** | **1 hr** | R-CS6C | Stimulated | 137.81 | 21.31 | 0.55 | 0.06 | 0.66 | 0.41 |
| | | | control | 121.9 | 28.01 | | | | |
| | | D-CS6C | Stimulated | 654.34 | 143.06 | 0.53 | 0.04 | 0.43 | 0.48 |
| | | | control | 524.37 | 59.44 | | | | |
| | **2 hr** | R-CS6C | Stimulated | 466.63 | 92.65 | 0.47 | 0.05 | 0.95 | 0.62 |
| | | | control | 471.62 | 72.18 | | | | |
| | | D-CS6C | Stimulated | 909.18 | 136.34 | 0.48 | 0.02 | 0.72 | 0.68 |
| | | | control | 942.28 | 132.3 | | | | |
| **Motor CX L5** | **1 hr** | R-CS6C | Stimulated | 367.24 | 62.4 | 0.51 | 0.07 | 0.97 | 0.83 |
| | | | control | 372.53 | 85.1 | | | | |
| | | D-CS6C | Stimulated | 1064.82 | 73.75 | 0.52 | 0.04 | 0.57 | 0.58 |
| | | | control | 1061.29 | 97.47 | | | | |
| | **2 hr** | R-CS6C | Stimulated | 502.76 | 92.70 | 0.49 | 0.03 | 0.77 | 0.85 |
| | | | control | 487.05 | 89.6 | | | | |
| | | D-CS6C | Stimulated | 912.71 | 136.35 | 0.49 | 0.009 | 0.68 | 0.83 |
| | | | control | 927.93 | 154.24 | | | | |

**Table S4. Numerical densities (NDs) of CS6C increase in response to sensory stimulation.** In layer 5 of the BCx, NDs of R-CS6C are increased at 1 hour survival time (i.e. sacrificed 1 hour following the end of passive whisker stimulation). D-CS6C NDs are not altered at this time point. Conversely, at 2 hours survival time, NDs of R-CS6Cs are not altered, but NDs of D-CS6C are significantly increased. To test for cortical area- and layer- specificity of this effect, R- and D-CS6Cs were counted in layer 5 of the motor cortex and layer 6 of the BCx. No changes were detected in either region/layer.

| Spine Type | Cluster Type | Mean | SEM | p value (2-way ANOVA) |
|---|---|---|---|---|
| **Mushroom Spines** | D-CS6C | 0.48 | 0.04 | 0.64 |
| | R-CS6C | 0.50 | 0.07 | |
| | Outside CS6C | 0.54 | 0.05 | |
| **Stubby Spines** | D-CS6C | 0.29 | 0.03 | 0.59 |
| | R-CS6C | 0.3 | 0.009 | |
| | Outside CS6C | 0.25 | 0.04 | |
| **Thin Spines** | D-CS6C | 0.17 | 0.02 | 0.15 |
| | R-CS6C | 0.14 | 0.03 | |
| | Outside CS6C | 0.28 | 0.05 | |

**Table S5.** Densities of dendritic spines morphologically identified as 'mushroom', 'stubby' and 'thin' (see Fig. S10) are not significantly different on dendrites embedded within D- or R- CS6Cs (n=5/CS6C type) and dendrites not associated with CS6Cs (n=5). Measurements were taken in n=5 mice.

| CS6C type | Spine measure | Location *(within/outside CS6C)* | Mean + SEM (μm) | Kolmogorov-Smirnov test | Mann-Whitney test |
|---|---|---|---|---|---|
| **R-CS6C** | Head Width | Within | 0.56 + 0.01 | **0.0001** | **0.00001** |
| | | Outside | 0.45 + 0.01 | | |
| | Head length | Within | 0.56 + 0.01 | **0.02** | **0.004** |
| | | Outside | 0.61 + 0.01 | | |
| | Neck length | Within | 0.44 + 0.01 | 0.69 | 0.34 |
| | | Outside | 0.45 + 0.01 | | |
| **D-CS6C** | Head Width | Within | 0.57 + 0.01 | **0.02** | **0.02** |
| | | Outside | 0.61 + 0.01 | | |
| | Head length | Within | 0.62 + 0.01 | **0.01** | **0.03** |
| | | Outside | 0.66 + 0.01 | | |
| | Neck length | Within | 0.44 + 0.01 | 0.08 | **0.02** |
| | | Outside | 0.51 + 0.01 | | |

**Table S6. Dendritic spine geometry is altered in R- and D- CS6Cs.** Within R-CS6Cs, mushroom spines have wider and shorter head, but unaltered neck length (n=194 spines over 4 dendrites) with respect to dendritic spines located on dendrites in the immediate vicinity, but outside, R-CS6Cs (n=154 spines over 4 dendrites). Wider and shorter spine heads within R-CS6C are interpreted to reflect synaptic potentiation (*2*). In contrast, within D-CS6Cs, mushroom spines have narrower and shorter head width and neck length (n=217 spines over 6 dendrites) with respect to dendritic spines located on dendrites outside R-CS6Cs (n=206 spines over 7 dendrites). The narrower head width potentially reflects synaptic de-potentiation (*2*).

**Supplemental Figures**

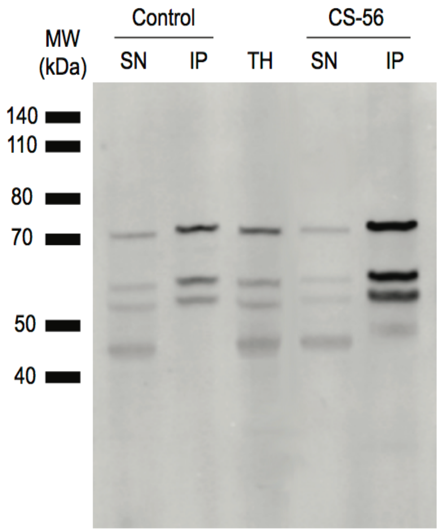

**Fig. S1 - CS-56 Immunoprecipitation**
Immunoblot of CS-56 immunoprecipitation from human postmortem amygdala tissue using mouse anti-CS-56 antibody. CS-56 was enriched in multiple protein bands in the IP. 20 µg of amygdala total homogenate was loaded as a positive control. Mouse IgM was used as a negative control. IP, immunoprecipitation; SN, supernatant; TH, total homogenate; MW molecular weight.

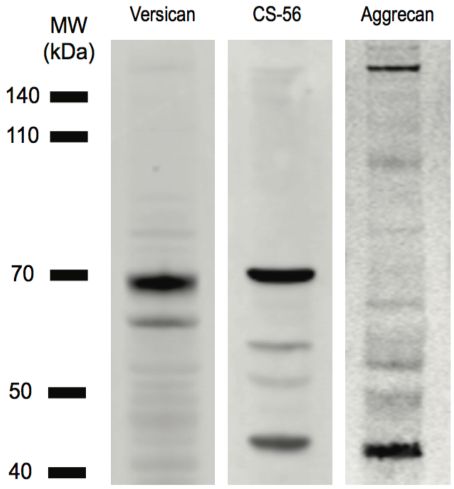

**Fig. S2 - Immunoblot study confirms presence of versican, CS-56 and aggrecan previously identified by CS-56 IP-MS within the human in amygdala**.
20 µg of total homogenate was probed with rabbit anti-versican, mouse anti-CS-56 and rabbit anti-aggrecan respectively. Protein bands were detected at comparable MW to those seen in CS-56 IP (Fig. S1). IP, immunoprecipitation; MW, molecular weight.

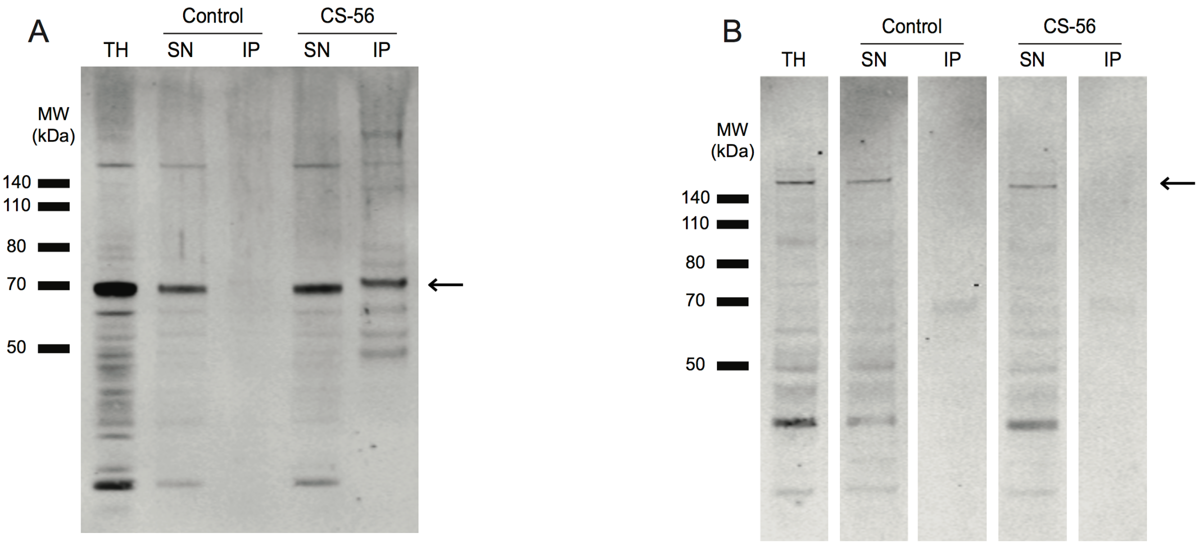

**Fig. S3 - Versican and Aggrecan Expression in CS-56 Immunoprecipitation**. ***A)*** Immunoblot showing expression of versican, the most abundant chondroitin sulphate proteoglycan protein detected in the CS-56 IP as determined by mass spectrometry. Versican protein fragments (band identified by arrow) were detected using rabbit anti-versican antibody following affinity purification of CS-56 in postmortem amygdala. 20 µg of amygdala total homogenate (TH) was loaded as a positive control. Mouse IgM was used as a negative control. ***B)*** Immunoblot showing expression of aggrecan, in supernatant but not eluate, following CS-56 affinity purification (identified by arrow). Aggrecan is a member of the chondroitin sulphate proteoglycan family of proteins but was not detected following mass spectrometry of the immunoprecipitate. Aggrecan acts as an experimental control, suggesting that those proteins identified by CS-56 IP-MS form a network of proteins that are specific to the CS-56 complex. 20 µg of amygdala total homogenate was loaded as a positive control. Rabbit IgG was used as a negative control. The lanes in Figure S3B were reordered during figure production to allow for comparison between.
*Abbreviations: IP, immunoprecipitation; SN, supernatant; TH, total homogenate; MW molecular weight.*

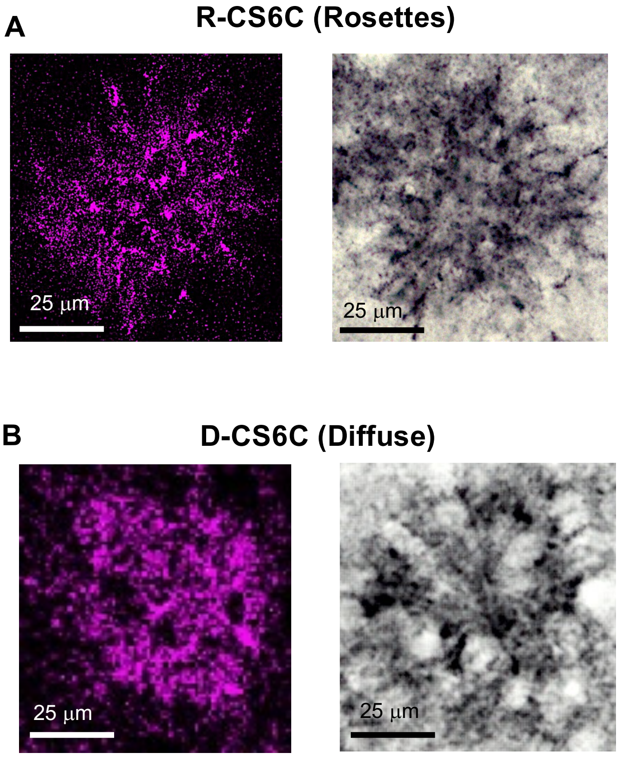

**Fig. S4 - Examples of CS6Cs in the mouse Barrel Cortex**
CS6Cs in the mouse BCx, immunolabeled with CS56 antibody and visualized with fluorescent (left) or light microscopy (right). The diameter of CS6Cs in mice is typically 75-100 µm. CS6Cs appear along a spectrum of morphological configurations. For the purpose of this study, they were classified in two main types, R-CS6Cs (A) and D-CS6Cs (B), according to their predominant features. Intermediate types were included in the D-CS6C group for quantification purposes. (A) R-CS6Cs: The predominant feature of this CS6C type is a multitude of intensely immunostained convergent processes. (B) D-CS6Cs: These CS6Cs present with the same shape and size as R-CS6Cs but are composed predominantly of intensely immunolabeled puncta.

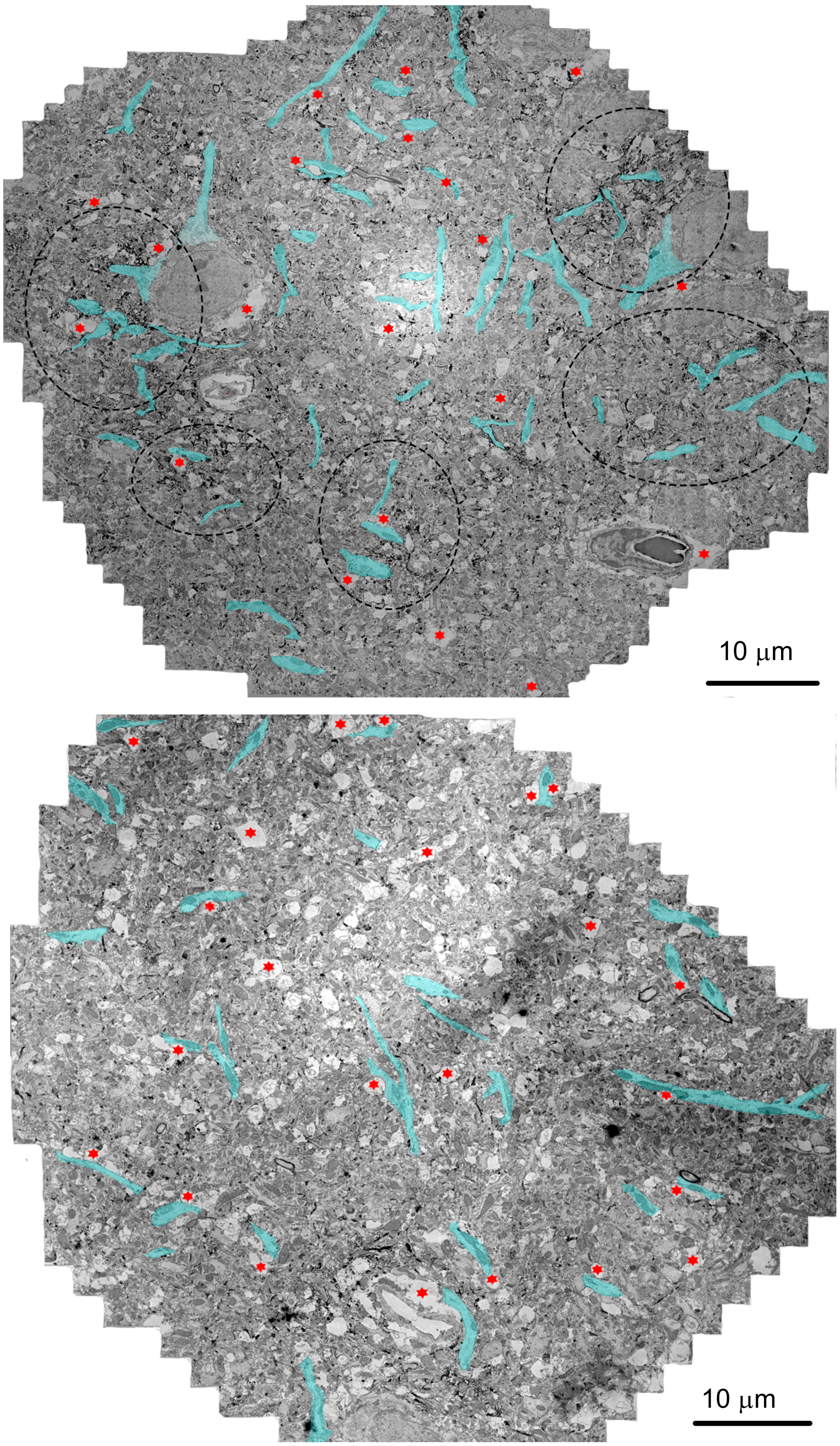

**Fig. S5. CS6Cs contain a multitude of dendrites and associated glial processes.** Electron microscopy photocomposites of a R-CS6C (top) and a D-C6C (bottom). Dendrites (pseudo-colored in blue) are in close contact with glial processes (red stars) and CS6 immunoprecipitate (black). In the top panel, areas of intense CS6 immunolabeling are marked by dashed lines.

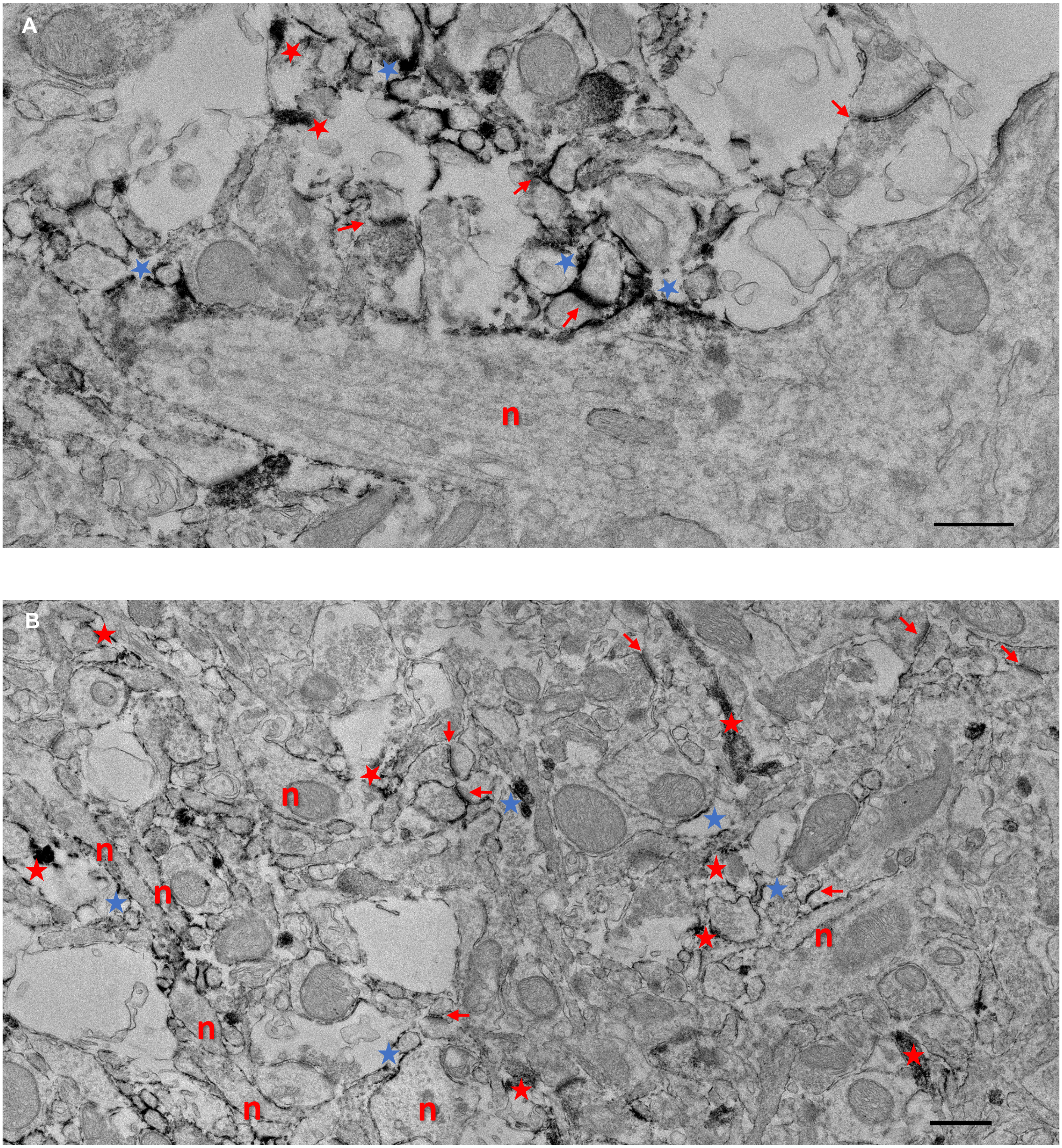

**Fig. S6 - CS6C ultrastructural morphology in the mouse Barrel Cortex (BCx).** Electron microscopy studies on BCx tissue investigated the ultrastructural characteristics of CS6Cs. Immuno-electron micrographs within a R-CS6C **(A)** and a D-CS6 **(B)** showing prominent immunoperoxidase reaction product within glial processes (red stars) and end-feet (blue stars), as well as within the synaptic cleft and around synaptic elements (red arrows). Note that some immunolabeling can also be detected on the outer surface of a dendritic plasma membrane, while the dendrite cytoplasm is void of labeling (Scale bar: 500nm

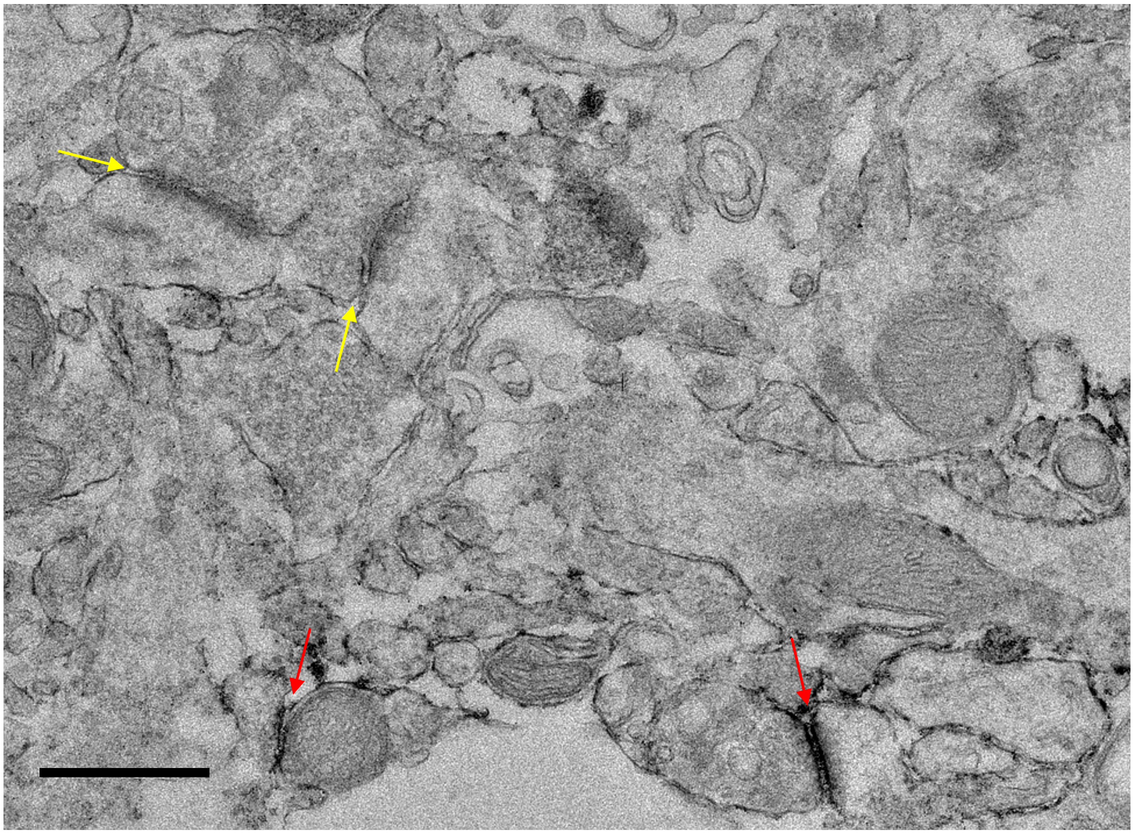

**Fig. S7 - Control for CS6C ultrastructural morphology in the mouse Barrel Cortex (BCx).** As a control for the accuracy of the identification of CS56-IR synapses, we tested whether CS56 immuno-negative synapses could also be detected with CS6Cs. Immuno-electron micrograph within a “R-CS6C” showing CS56-immunolabeled synapses (red arrows) in close proximity of immuno-negative synapses (yellow arrows) (Scale bar: 500 nm).

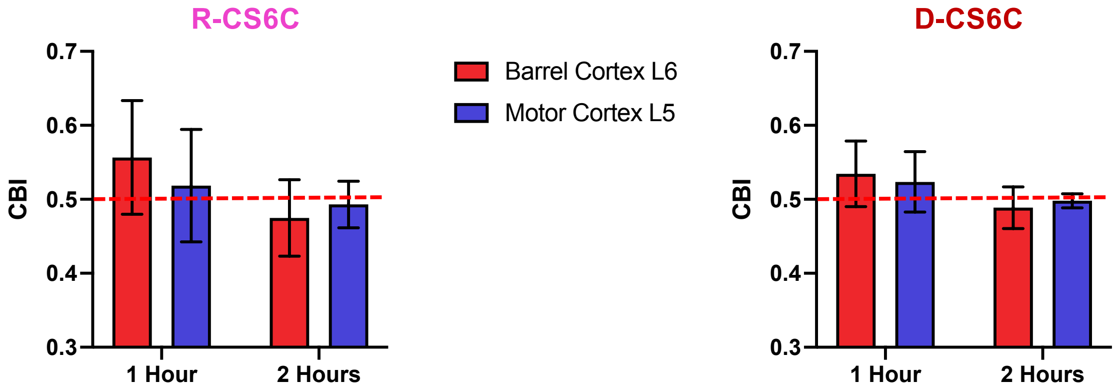

**Fig.S8 - Sensory manipulation experiments – Additional data and control studies A)** CS6C numerical densities (NDs) in the BCx do not differ between hemispheres in control animals (p=0.55), but are significantly decreased in the deprived hemisphere of sensory-deprived animals with respect to the control hemisphere (p=0.01). **B)** Contralateral bias index (CBI) analysis in control mice shows comparable CS6Cs-NDs between hemispheres across layers. (L2/3 p=0.34. L4 p=0.26. L5 p=0.71. L6 p=0.92) **C)** Representative picture depicting reduced CS6Cs NDs in a sensory-deprived hemisphere (bottom left panel) compared to a not-deprived (bottom right panel)

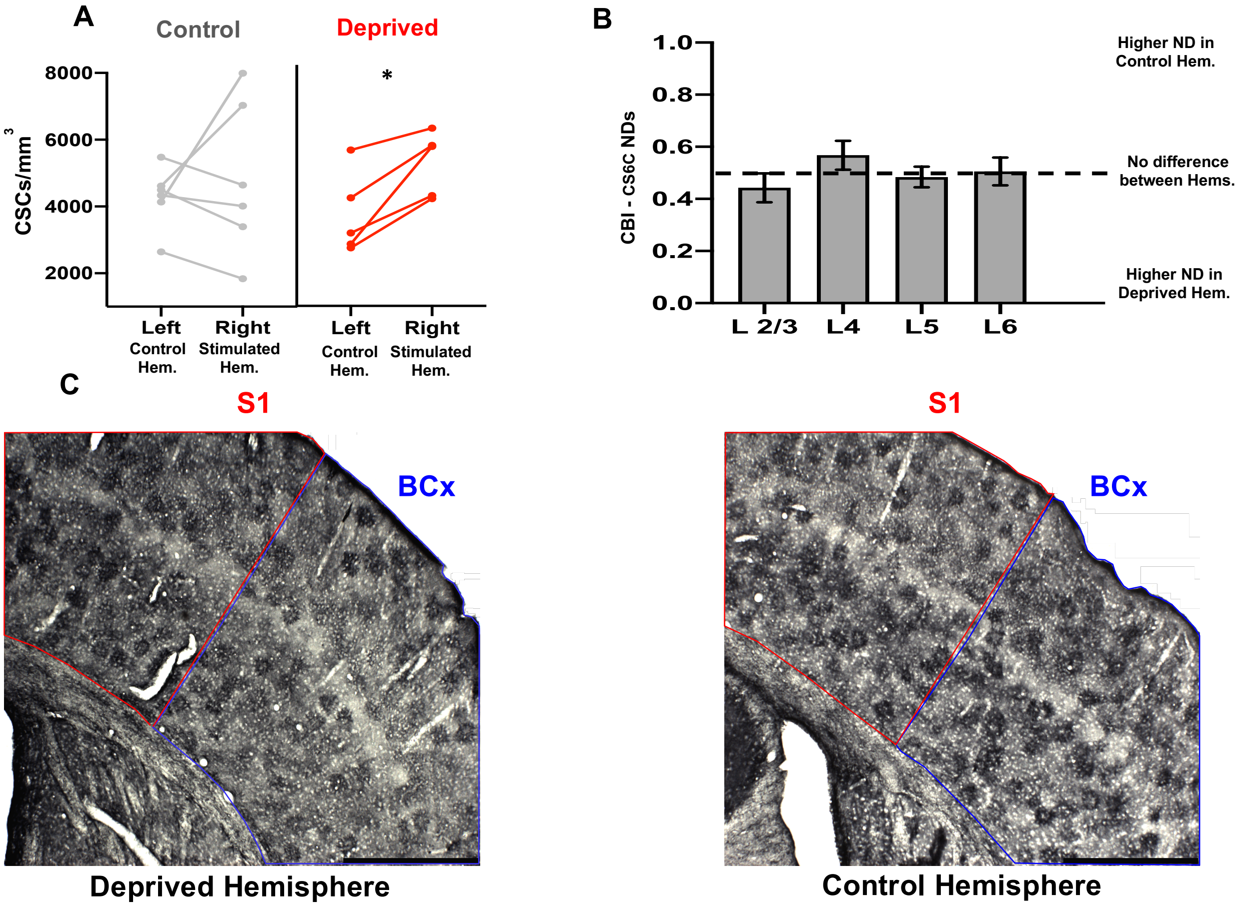

**Fig. S9 - Control brain regions, i.e. layer 5 of the Motor Cortex and Layer 6 of the BCx, do not show significant differences of CS6C NDs.** CBI analysis at two different time points (1 hr and 2 hr) and for R-CS6Cs and D-CS6C shows that NDs are not significantly different between the stimulated and control hemisphere.

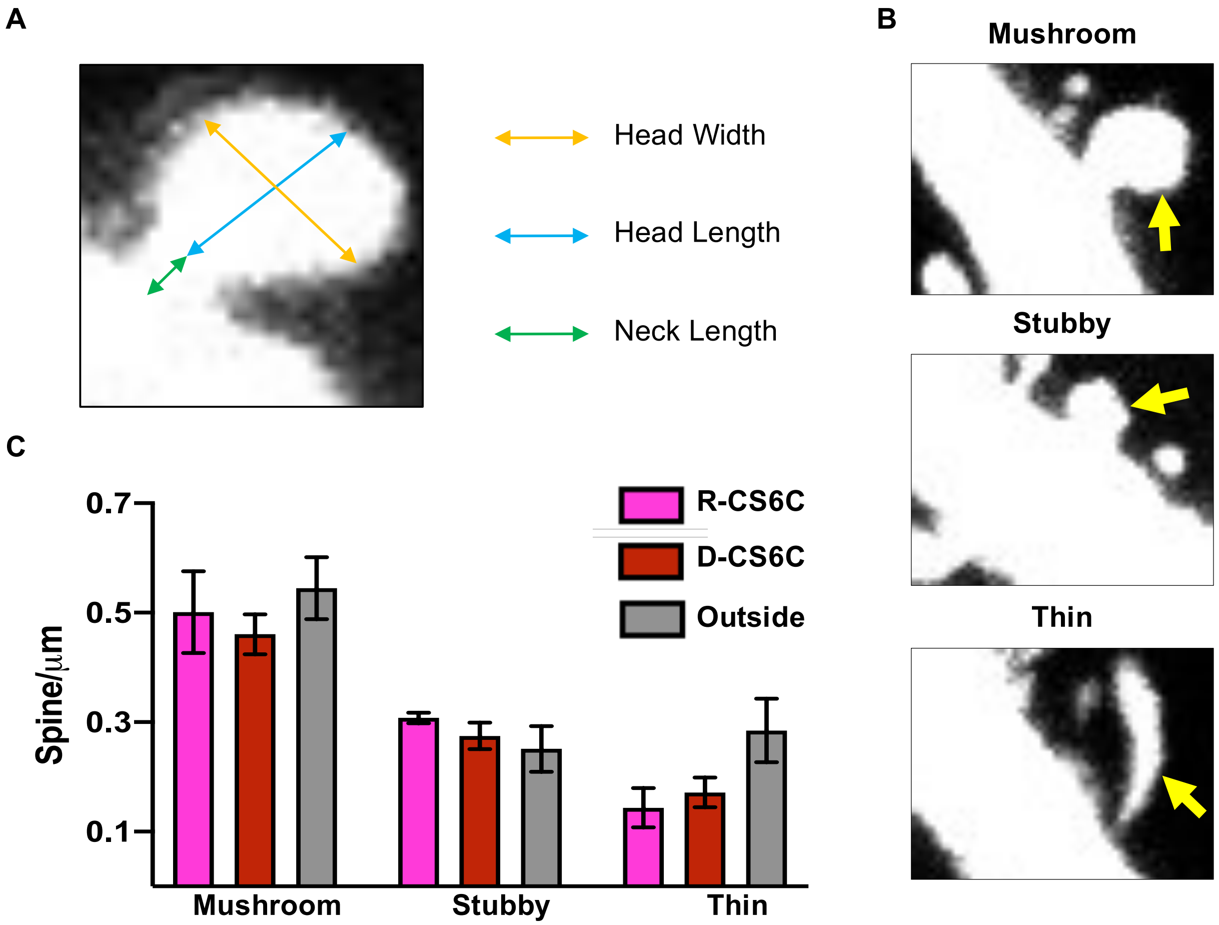

**Fig. S10 Dendritic spines' analyses.** As a first step toward testing the hypothesis that CS6Cs are associated with structural plastic changes at synapses we assessed the dendritic spines' density within and outside CS6Cs. **A)** Representative picture showing the method for dendritic spine's measurement. Total spine length was obtained combining neck and head length. **B)** Confocal photomicrographs showing examples of mushroom, stubby and thin dendritic spines spines classification (yellow arrow). Classification of dendritic spines was performed according to the following criteria. *Mushroom* spines had clearly distinguishable spine head and neck, with the neck diameter significantly narrower than the head diameter. *Stubby* spines did not have a clearly distinct neck. *Thin* spines had a long, thin necks and heads and the total length of the spine was considerably greater than spine width. **C)** Dendritic spine density for mushroom, stubby and thin spine is not different within CS6Cs with respect to areas outside CS6Cs.

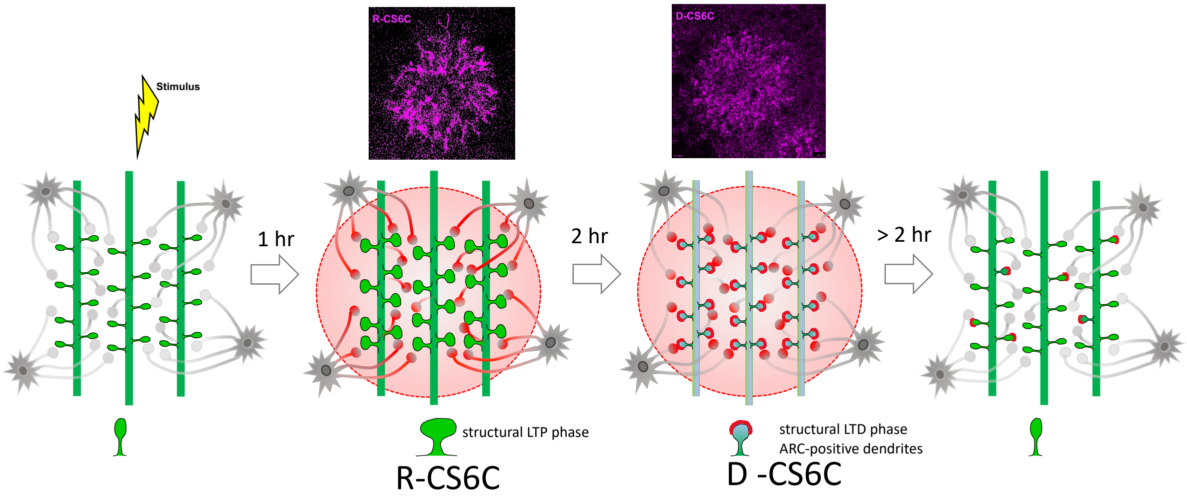

**Fig. S11. Proposed model for CS6C (pale red circles) as dynamic segregated microenvironments involved in coordinated synaptic plasticity across multiple neighboring dendrites.** Within 1 hour following stimulation, CS6 moiety (red) is transported from glial cells through their processes (R-CS6Cs), converging onto a population of neighboring synapses. R-CS6Cs transition into D-CS6Cs once CS6 is secreted into the synaptic cleft and surrounds postsynaptic terminals. This transition corresponds to increased dendrite ARC expression within the D-CS6Cs and a synaptic structural LTD phase. Following this phase, only sparse CS6-coated synapses may be present and CS6Cs can no longer be detected
